# Detecting and correcting for bias in Mendelian randomization analyses using gene-by-environment interactions

**DOI:** 10.1101/187849

**Authors:** Wes Spiller, David Slichter, Jack Bowden, George Davey Smith

## Abstract

**Background:** Mendelian randomization has developed into an established method for strengthening causal inference and estimating causal effects, largely due to the proliferation of genome-wide association studies. However, genetic instruments remain controversial as pleiotropic effects can introduce bias into causal estimates. Recent work has highlighted the potential of gene-environment interactions in detecting and correcting for pleiotropic bias in Mendelian randomization analyses.

**Methods:** We introduce MR using Gene-by-Environment interactions (MRGxE) as a framework capable of identifying and correcting for pleiotropic bias, drawing upon developments in econometrics and epidemiology. If an instrument-covariate interaction induces variation in the association between a genetic instrument and exposure, it is possible to identify and correct for pleiotropic effects. The interpretation of MRGxE is similar to conventional summary Mendelian randomization approaches, with a particular advantage of MRGxE being the ability to assess the validity of an individual instrument.

**Results:** We investigate the effect of BMI upon systolic blood pressure (SBP) using data from the UK Biobank and the GIANT consortium using a single instrument (a weighted allelic score). We find MRGxE produces findings in agreement with MR Egger regression in a two-sample summary MR setting, however, association estimates obtained across all methods differ considerably when excluding related participants or individuals of non-European ancestry. This could be a consequence of selection bias, though there is also potential for introducing bias by using a mixed ancestry population. Further, we assess the performance of MRGxE with respect to identifying and correcting for horizontal pleiotropy in a simulation setting, highlighting the utility of the approach even when the MRGxE assumptions are violated.

**Conclusions:** By utilising instrument-covariate interactions within a linear regression framework, it is possible to identify and correct for pleiotropic bias, provided the average magnitude of pleiotropy is constant across interaction covariate subgroups.

Key Messages

- Instrument-covariate interactions can be used to identify pleiotropic bias in Mendelian randomization analyses, provided they induce sufficient variation in the association between the genetic instrument and exposure.
- By regressing the gene-outcome association upon the gene-exposure association across interaction covariate subgroups, it is possible to obtain an estimate of the average pleiotropic effect and a causal effect estimate.
- The interpretation of MRGxE is analogous to that of MR-Egger regression.
- The approach serves as a valuable test for directional pleiotropy and can be used to inform instrument selection.

## Introduction

Mendelian randomization (MR) has developed into a popular multifaceted approach to strengthening causal inference in epidemiology(1, 2). In many cases, MR analyses involve employing genetic variants as instrumental variables (IVs) allowing for causal effects to be consistently estimated in the presence of unmeasured confounding. This requires candidate variants to be associated with the exposure of interest (IV1), not be associated with confounders of the exposure and outcome (IV2), and not be associated with the outcome through pathways outside of the exposure (IV3)(3). The extent to which specific genetic variants satisfy these assumptions is often controversial, however, due to uncertainties around the true mechanisms responsible for observed gene-phenotype relationships(4).

One issue of particular concern is potential violation of IV3 through horizontal pleiotropy-occurring when a genetic instrument is associated with a study outcome through biological pathways outside the exposure of interest(2, 5). This introduces bias into causal effect estimates in the direction of the pleiotropic association, and can inflate type I error rates when testing causal null hypotheses(5, 6). In an individual level data setting it is common practice to combine genetic variants into allelic scores, creating a single stronger instrument. In ideal cases, an instrument which is sufficiently strong relative to the average pleiotropic effect across all the constituent variants may potentially mitigate bias due to pleiotropy, however, it is possible that such an approach can exacerbate bias by obscuring inconsistencies in genetic associations(7, 8). Where multiple instruments are available, an alternative strategy is to adopt a meta-analytic approach(9). If the set of genetic variants do not exhibit an average non-zero (or ‘directional’) pleiotropic effect an inverse variance weighted (IVW) estimate can be used to obtain an effect estimate equivalent to two-stage least squares (TSLS) regression(9). In cases where directional pleiotropy is suspected, MR-Egger regression can be used to estimate and correct for pleiotropic bias provided the variants’ pleiotropic effects are independent of their strengths as instruments (the InSIDE assumption)(5). It is also possible to implement weighted median and mode-based estimation to quantify the magnitude of pleiotropic effects, and provide a corrected causal effect estimate(10, 11). Such methods are most applicable in two-sample summary MR(12, 13).

In the econometrics literature, Slichter regression has been proposed as a method for evaluating instrument validity within a potential outcomes framework(14, 15). This involves either observing or extrapolating to a population subgroup for which the instrument and exposure are independent (defined as a *no relevance group*), and then measuring the association between the instrument and outcome which would arise for such subgroups. In cases where such a subgroup with adequate power to detect an association is not present, the association for a hypothetical no relevance group can be estimated. A non-zero instrument-outcome association in such a sub-group serves as evidence that IV3 has been violated. Slichter regression builds upon a number of key developments in econometrics, in particular the identification and estimation of local average treatment effects put forward by Imbens and Angrist(15). Works such as Card(16) have focused upon heterogeneity analysis utilising geographic, age, ethnicity, and family background characteristics to estimate causal effects; such returns to schooling by considering an observed interaction between college proximity and IQ. Conley et al(17) emphasise the potential trade-off between instrument strength and degree of IV3 violation in putting forward the notion of plausibly endogeneity, whilst further works underlining the utility of using instrument-covariate interactions have also emerged, such as those of Gennetian et al and Small(18, 19). It is also worth noting that a similar method, entitled Pleiotropy Robust Mendelian Randomization (PRMR), has recently been put forward with a similar intuitive framework(20). However, the PRMR approach requires a subgroup for which the instrument and the exposure are independent to be observed, potentially limiting the applicability of the approach in contrast to approaches which estimate the expected association at a hypothetical independent subgroup.

In this paper, we introduce Slichter regression within the context of epidemiology, and in doing so formalise the increasing use of gene-environment interactions for assessing instrument validity(21–27). In doing so, we present MR using Gene-by-Environment interactions (MRGxE) as a statistical framework and sensitivity analysis to identify and correct for pleiotropic bias in MR studies using gene-covariate interactions. An important aspect of MRGxE is the ability to assess the validity of a single instrument, in contrast to methods examining heterogeneity across a set of MR estimates using many instruments.

Two key features differentiate MRGxE from analogous methods in the econometrics literature. Firstly, MRGxE is conducted within a linear regression framework as opposed to utilising local linear regression techniques commonly used in such approaches. This has the advantage of improving the ease with which MRGxE can be implemented and has similarities in interpretation to conventional two-sample summary MR sensitivity analyses. Additionally, MRGxE can be applied in cases where only summary data are available and is therefore not contingent upon the availability of individual level data. Such data could be obtained from previously published studies in cases where subgroup specific estimates are provided, as is increasingly common in the case of gender, or alternatively requested from consortia without requiring full access to individual level data.

We begin by outlining the MRGxE framework, highlighting the assumptions and implementation of the approach. The structure of MRGxE is similar to LD score regression(28), interpreting the intercept as a bias term within the underlying model, and can be viewed as an analogous approach to MR-Egger regression substituting population subgroups for genetic variants. This allows for estimation of the validity of individual instruments. With this complete, an applied example is considered examining the effect of body mass index (BMI) upon systolic blood pressure (SBP) using the most recent release of data from UK Biobank (July 2017) and the GIANT consortium(29). Initially, a two-sample summary MR analysis is conducted as a frame of reference, using derived summary statistics from each sample. With this complete, MRGxE is implemented using a single weighted allelic score. Using MRGxE without excluding participants on the basis of non-European ancestry or relatedness we find evidence suggesting a positive association between BMI and SBP. However, when utilising exclusion criteria, the findings are suggestive of instrument invalidity with limited indication of an association between BMI and SBP. This is most likely the result of selection bias induced by sub setting the data, as the contrasting effect estimates are also observed using two-sample summary MR approaches. Importantly, it is the observed agreement between MRGxE and two-sample summary approaches that is highlighted, as correcting for selection bias is beyond the scope of this work. We examine the results within the context of underlying assumptions of the model, conducting a simulation study to demonstrate the effectiveness of the approach under varying conditions.

## Methods

### Non-technical intuition

Consider a situation in which the instrument-exposure association is found to vary between subgroups of the target population. We follow Slichter(14) in defining an observed subgroup for which the instrument does not predict the exposure of interest as a *no relevance group*. As a valid genetic instrument can only be associated with the outcome of interest through the exposure, it follows that a valid instrument would also not be associated with the outcome for a no relevance group. Any non-zero instrument-outcome association for the no relevance group can therefore be interpreted as evidence of horizontal pleiotropy.

This intuitive approach to pleiotropy assessment has been considered in a number of epidemiological studies. For example Chen et al(26) considered differences in drinking behaviour by gender in East Asian populations within a fixed effects meta-analysis of the *ALDH2* genetic variant and blood pressure. Observing that males are much more likely to consume alcohol than females, gender-stratified drinking behaviour was used to identify female participants as a no relevance group. This interaction has also received further attention in work such as Cho et al(21) and Taylor et al(30).

Previous applications considering variation in instrument-exposure association across populations extend beyond simple gender differences, such as Tyrrell et al(23) investigating the extent to which genetically predicted BMI is associated with environmental factors through gene-covariate interactions. They identified genetically predicted BMI as a weaker instrument for participants experiencing lower levels of socio-economic deprivation (as quantified by the Townsend deprivation index), and utilised negative controls to examine residual confounding(23). A further interesting example is stratifying by smoking status, as considered in Freathy et al(31) in their examination of the relationship between genetic instruments used to predict smoking status and adiposity. In their recent work, Robinson et al(32) identify genotype-covariate interactions with respect to the heritability of adult BMI, finding evidence of genotype-age and genotype-smoking interactions. Gene-environment interactions with covariates such as socio-economic status were not identified as having a substantial impact on the distribution of phenotypic effects, though this may be the result of a lack of statistical power or measurement error in self-reported covariates.

In presenting MRGxE we highlight similarities to the approach of Cho et al(21), in which a gender-*ALDH2* interaction term was incorporated within a two-stage least squares (TSLS) model to estimate the degree of horizontal pleiotropy. In doing so, we clarify how it works when individual level data are available, and crucially demonstrate how MRGxE extends this approach so that it can be additionally applied to summary data. This extends the applicability of the method to two sample summary data MR, and also general meta-analysis contexts.

### The MRGxE Framework

Consider an MR study consisting of *N* participants (indexed by *i* = 1, …, *N*). For each participant, we record observations of a genetic instrument *G_i_* an exposure *X_i_*, an outcome *Y_t_*, and a further covariate *Z_i_* which induces variation the association between *G_i_* and *X_i_* through an interaction *GZ_i_*. The relationship between each variable is illustrated in Figure 1, with *U* representing a set of all unmeasured variables confounding *X* and *Y*, and *I_GZ_* representing the interaction term for ease of interpretation.

**Figure 1:**
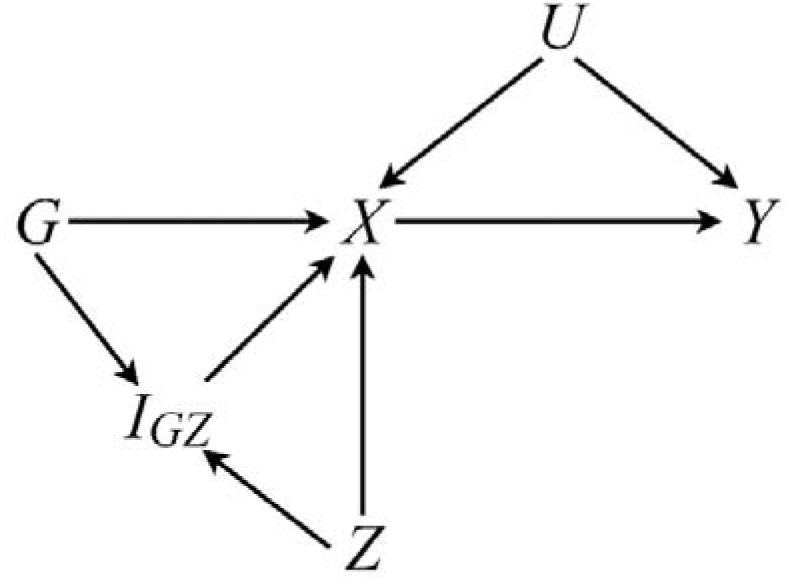
A directed acyclic graph (DAG) showing the assumed relationship between each variable in MRGxE.

The exposure *X* is considered a linear function of *G, Z, GZ, U* and an independent error term, *ϵ_x_*, whilst the outcome *Y* is a linear function of *G, Z, GZ, U, X* and an independent error term, *ϵ_Y_*. Using *γ* and *β* to denote regression coefficients for the first and second stage models respectively, a two-stage model can be defined as:

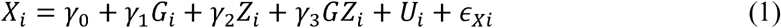

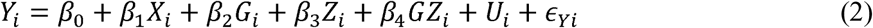

The causal effect of *X* on *Y* is denoted *β*_1_ and is the parameter we wish to estimate. The pleiotropic effect of the instrument across the sample is *β*_2_. Note that performing an ordinary least squares (OLS) regression of *Y* upon *X* would yield a biased estimate of *β*_1_ due to confounding, and a TSLS regression of *Y* on the genetically predicted exposure 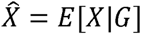 would result in biased estimates in cases where *β*_2_ ≠ 0. This is demonstrated through a simple simulation presented in the web appendix.

At its core, MRGxE utilises a gene-covariate interaction as an instrument. This subsequently places restrictions on the interaction analogous to the conventional relevance (IV1), exogeneity (IV2), and exclusion restriction (IV3) assumptions of IV analysis. A suitable interaction *GZ* is therefore:

1. Associated to the exposure of interest (*γ*_3_ ≠ 0).
2. Not associated with confounders of the exposure and outcome (*GZ* ⊥ *U*).
3. Not associated with the outcome outside of the exposure of interest (*β*_4_ = 0).

The first assumption can be assessed by directly fitting the first stage model. Regarding the second assumption, it is crucial to highlight that it pertains to the independence of the **interaction** with respect to confounders, and not *G* and *Z* individually. Finally, the third assumption requires pleiotropic effects remain constant across the population. Variation in pleiotropic effect is in many cases driven by violations of the second assumption, though we defer discussion of these mechanisms to the next section.

A value of *Z* defining a no relevance group (either observed or hypothetical) can be determined using model (1) by deriving the covariate value *Z = z_x_* at which *G* and *X* are independent. This is achieved by calculating the partial effect of *G* upon *X* and rearranging such that:

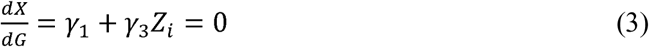

This yields the trivial solution

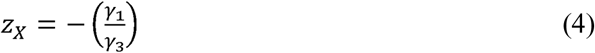

In cases where the covariate value *z_X_* is actually observed in the population, regressing *Y* upon *G* for the subset of participants with *Z* = *z_X_* will provide an estimate of horizontal pleiotropy (that is for *β*_2_) as the coefficient of *G*. Unfortunately, this approach is difficult to implement in practice, either because the value *z_X_* is not observed in the population or the subset of participants is simply too small to provide sufficient power. As a consequence, it is often more appropriate to estimate the degree of pleiotropy at a theoretical (or extrapolated) no-relevance group, using differences in instrument-exposure associations across values of *Z*.

To illustrate how this is possible, a *reduced form* IV model is constructed, that is, models for *X* given *G*, and *Y* given *G* by rewriting model (1) as

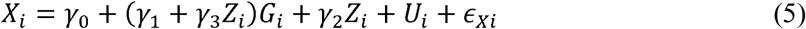

and model (2) as

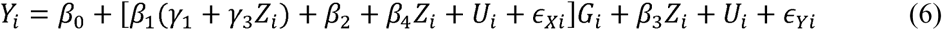

The change in *G* – *X* and *G* – *Y* associations for a given change in *Z* can be identified as the coefficient of *G* in models (5) and (6) respectively (with *β*_4_ set to 0) as

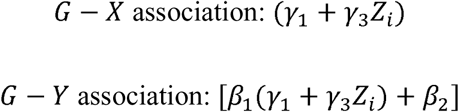

The Wald ratio(33) estimand for the causal effect of *X* on *Y* (i.e. the true *G* – *Y* association divided by the true *G* – *X* association) would then be equal to:

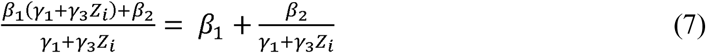

That is, the causal effect, *β*_1_ plus a non-zero bias term whenever *β*_2_ is non-zero. In the Cho et al(21) analysis, an estimate for *β*_1_ was obtained by performing TSLS regression using the interaction as the instrument, by fitting models (8) and (9) below:

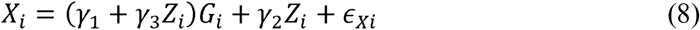

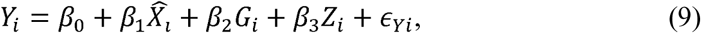

Where 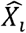 is the fitted value from model (8). In this case, the coefficient *β*_2_ represents the degree of pleiotropy for the genetic instrument *G*. Cho et al also demonstrate the use of two-stage predictor substitution (TSPS) models of the same structure when considering binary outcome variables(21).

Whilst this approach is useful in not requiring an observed no relevance group, it has two limitations. First, as a consequence of utilising TSLS and TSPS, it is only applicable to cases in which individual level data are available. In utilising genetic data, it has become common to utilise summary data within a meta-analysis context, as individual studies often lack statistical power due to sample size restrictions. A second limitation is that it assumes an underlying linear model, which may not hold in practice. For example, in considering adiposity as an exposure, individuals at extreme values could be at greater risk, implying a curved relationship, but this fact would not be apparent from fitting the above models. Care is therefore needed in investigating and justifying the assumption of an underlying linear model. MRGxE attempts to overcome these limitations by reframing the model within a two-sample summary MR context, delivering a consistent estimate for *β*_1_ by executing the following three step procedure:

1. Estimate *G* – *X* and *G* – *Y* associations at a range of values of Z.
2. Regress the *G* – *Y* associations on the *G* – *X* associations within a linear regression.
3. Estimate the causal effect *β*_1_ as the slope of the regression.

Let *Z_j_* denote the *j^th^* subgroup of *Z* (*j* = 1,…,*J*). For each group *Z*_j_, we initially estimate the instrument-exposure association and standard error (step 1) by fitting the following regression model:

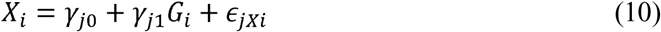

Note that we include a subscript *j* to distinguish the regression parameters from the first stage model (1). The coefficient *γ*_*j*1_ is therefore interpreted as the *G* – *X* association for group *Z_j_*. Next, we fit the corresponding instrument-outcome regression model (step 2):

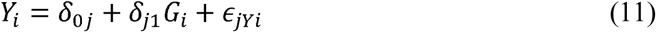

In this case, we use *δ*_*j*1_ to denote the *G* – *Y* association coefficient for group *Z_j_*, distinguishing the model from (6). Thus, from models (10) and (11) we obtain sets of *G* – *X* associations (*γ*_*J*1_) and *G* – *Y* associations (*δ*_*J*1_) respectively across *Z_J_* subgroups. Finally, we regress the set of 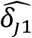 estimates upon the set of 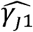 estimates (step 3):

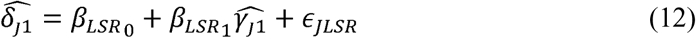

In Model (12), 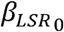 is an estimate of directional pleiotropy (*β*_2_) whilst 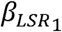 is the causal effect of *X* upon *Y* corrected for any directional pleiotropy (*β*_1_) To understand how this is the case, recall that *β*_2_ represents a constant pleiotropic effect across subgroups of *Z*. Model (12) can be thought of as an average of the ratio estimates across *Z_J_*, with the bias parameter *β*_2_ estimated as the intercept. This is illustrated in Figure 2.

**Figure 2:**
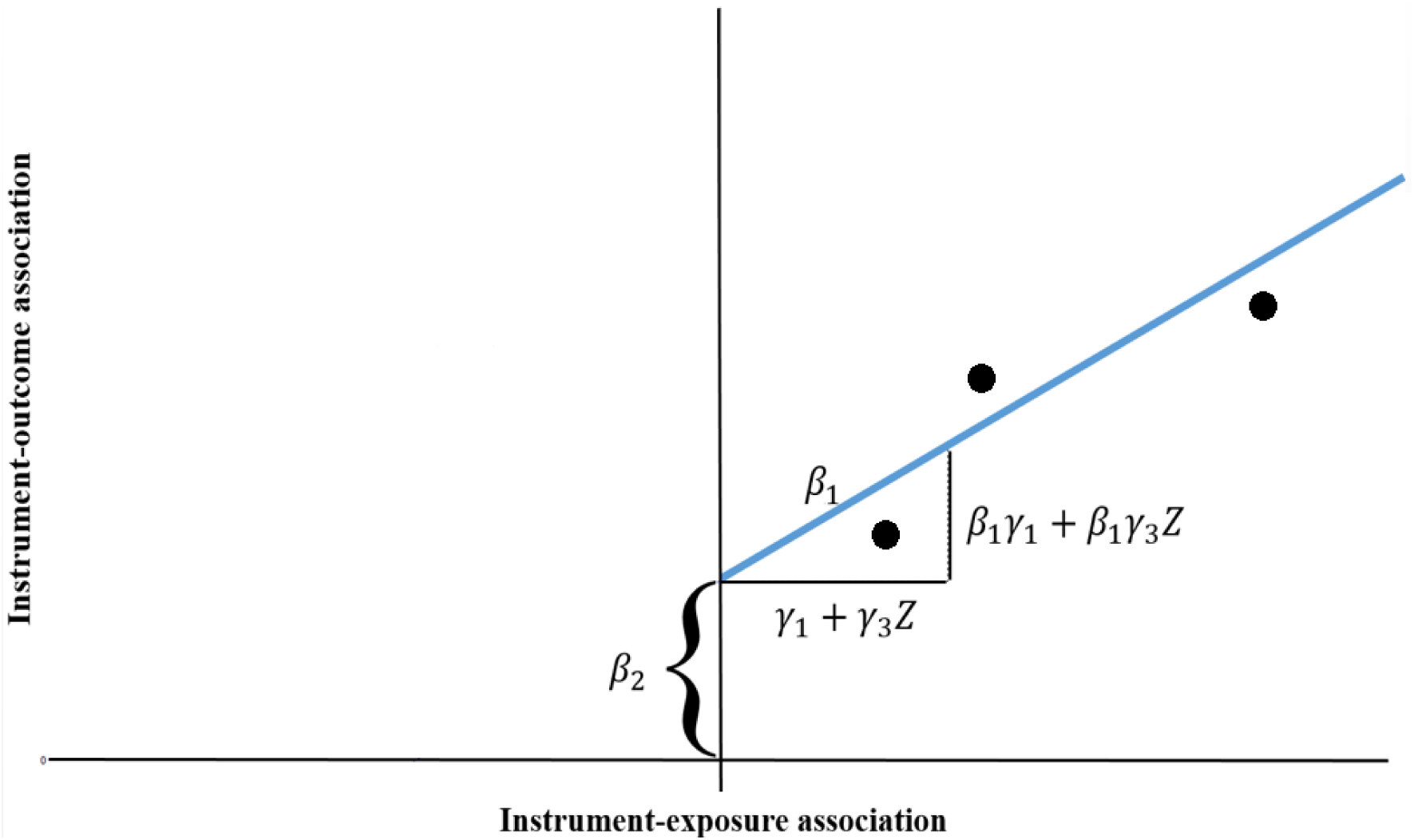
Hypothetical plot showing components of MRGxE. For a set of groups represented as solid points, the x-axis represents the association between the genetic instrument and the exposure, whilst the y-axis shows the association between the genetic instrument and the outcome. The point at which is an estimate of the theoretical no relevance group, with a remaining (pleiotropic) association between the instrument and the outcome given as the intercept ( ).

Provided an appropriate set of subgroups is defined, one advantage of producing a plot as shown in Figure 2 is that the functional form of the interaction can potentially be inferred from the distribution of subgroups. This is demonstrated within a simulation provided in the web appendix, using linear, quadratic, and cubic interactions. To show how the intercept estimates *β*_2_, consider the reduced form model (6) evaluated for the no relevance group *Z* = *z_X_*. From equation (4), 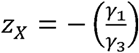. Then, by substitution:

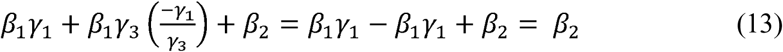

In cases where the intercept estimate passes exactly through the origin, the MRGxE causal effect estimate would be identical to the inverse-variance weighted (IVW) estimate. This mirrors the equivalence of IVW and MR-Egger regression in the multiple instrument setting when the estimated average pleiotropic effect across all variants is equal to 0. MRGxE can therefore be viewed as directly analogous to MR-Egger regression: in MRGxE we simply replace *G* – *X* association estimates for multiple variants with subgroup-specific associations for a single variant. R code for implementing MRGxE is provided in the web appendix.

At this point we highlight several important factors to consider when implementing MRGxE. Initially, it is important to define a sufficient number of subgroups of *Z* so as to accurately characterise the underlying gene-covariate interaction. Defining too few groups reduces power to detect either an association or instrument invalidity, due to the loss of information when combining two or more distinct groups. Where this is the case an estimate 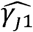 represents an average across the two groups, diminishing differences in instrument-exposure association across the set of estimates. Additionally, the lack of observations for instrument-exposure and instrument-outcome associations has an adverse effect on statistical power. The primary concern in selecting too many groups is that as the number of groups increases, the number of observations within each group decreases, reducing the precision of instrument-exposure and instrument-outcome association estimates. These features of group selection are explored in a simulation setting detailed in the web appendix.

Whilst MRGxE may appear most suited to settings for which we have a categorical interaction covariate, this is not necessarily the case. In many situations, categorical variables simply use predefined groups, whilst implicitly measuring a continuous covariate. For example, frequency of alcohol consumption focuses upon quantity of alcohol consumed over time, which is essentially continuous. We therefore suggest researchers to be critical of subgrouping imposed by using categorical variables where continuous data are available.

A second important consideration when performing MRGxE is to *not* transform effects to be positive using MRGxE as is advised for MR-Egger regression, as this mischaracterises the interaction term, attenuating causal effect estimates. A simulated example illustrating this issue is presented in the web appendix.

Finally, it is important to emphasise that in cases where instrument-exposure associations are present for all groups in the same direction, the accuracy in extrapolating the regression line towards a theoretical no-relevance group will be a function of the distance from the minimum *Z_j_* instrument-exposure association, and variation in the set of *Z_j_* instrument-exposure associations. This feature of MRGxE is examined further in the web appendix.

### The constant pleiotropy assumption

As a single (constant) parameter, *β*_2_ equates to the ‘correct’ intercept for MRGxE. That is the intercept which must be estimated in order to identify the correct causal effect, *β*_1_. Consistent estimates for both *β*_2_ and *β*_1_ are produced in cases where *β*_4_ = 0, that is, when the pleiotropic effect remains constant across all values of *Z*. If *β*_4_ ≠ 0 then the constant pleiotropy assumption is violated, which in our model would lead to the true pleiotropic effect *β*_2_ being equated to 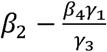. This would, in turn, lead to bias in the causal estimate for *β*_1_ such that:

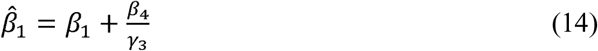

The derivation of this result is provided in the web appendix. From equation (14) it is clearly possible to mitigate the effect of bias due to violation of the constant pleiotropy assumption when the instrument-covariate interaction (*γ*_3_) is sufficiently large relative to the variation in pleiotropic effect *β*_4_, as the bias will tend towards zero *γ*_3_ as increases. However, as it is not possible to directly estimate *β*_4_ justifying the relative effect sizes of the first and second stage interactions would likely rely upon a priori knowledge.

Violations of the constant pleiotropy assumption can result from specific confounding structures in the underlying true model. Specifically, there must be no pathway from *G* to *Z* through the confounder *U*, no pathway from *Z* to *G* through *U*, and *U* cannot be a joint determinant of *G* and *Z*. Figure 3 shows a set of four possible scenarios in which the instrument *G* and interaction covariate *Z* are associated with a confounder *U*.

**Figure 3:**
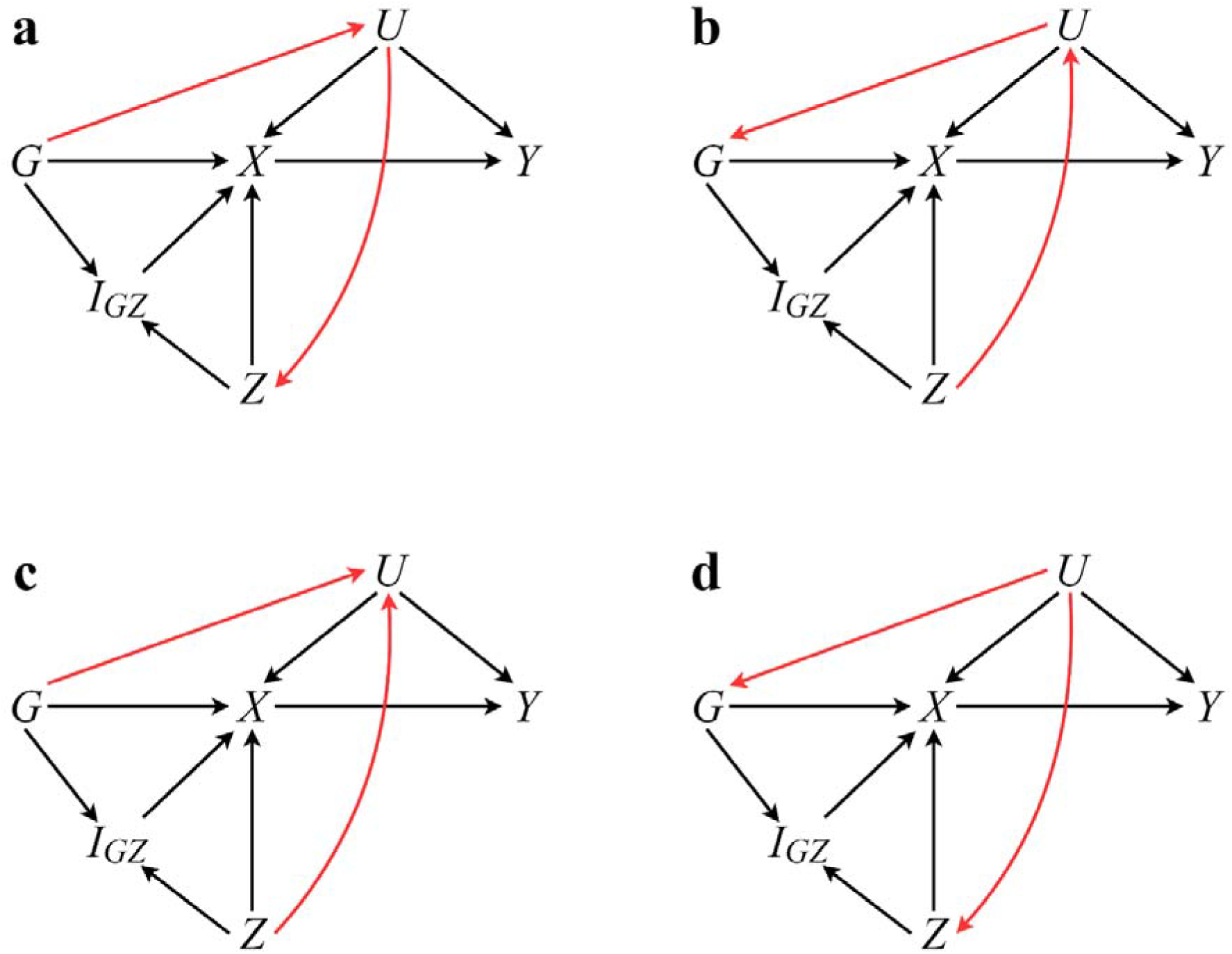
A set of DAGs illustrating confounding structures (indicated by red arrows) which invalidate the MRGxE approach. In this case, scenarios (a) (b) and (d) induce bias in MRGxE estimates. However, this is not the case for scenario (c) or when the confounder is associated with either or individually.

In Figure 3, scenarios (a) (b) and (d) introduce bias in MRGxE estimates, whilst scenario (c) and individual associations between either *Z* and *G* with *U* are not problematic. To illustrate how this is the case, we present a derivation of the MRGxE estimate for scenario (a) below, and present derivations for the alternate scenarios in the web appendix.

A set of data generating models mirroring scenario (a) are defined as

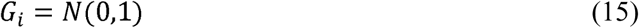

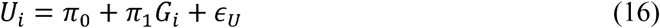

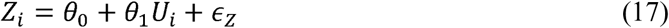

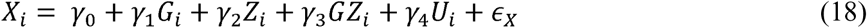

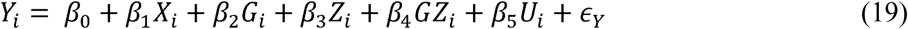

By substituting models (15), (16), and (17) into model (18) we can construct a model for *X* given *G* as

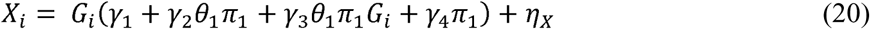

Where *η_x_* represents the combined error and intercept terms not pertaining to *G_i_*.

Using the same approach, we can also use model (19) to construct a model for *Y* given *G* as

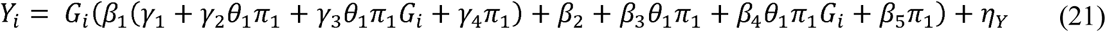

Where *η_Y_* represents the combined error and intercept terms not pertaining to *G_i_*. The corresponding Wald estimand using models (20) and (21) is then given as:

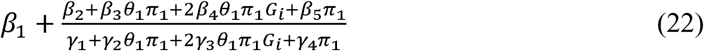

From equation (22) we can see that the constant pleiotropy assumption will be violated in cases where *θ*_1_ ≠ 0 and *π*_1_ ≠ 0, since the bias term is a function of the random variable *G*. This is the primary mechanism through which the constant pleiotropy assumption is violated. In the above, the parameter *β*_4_ represents the change in pleiotropic effect across the sample not captured by measured values *θ*_1_ and *π*_1_, though in many cases these will not be measured and can be folded into the *β*_4_ term. In scenario (b), the result is similar to equation (23) with the exception that *Z* is included instead of *G*, though importantly estimates using MRGxE will not exhibit bias where *θ*_1_ = 0 or *π*_1_ = 0.

As a consequence, the range of interaction covariates suitable for use within MRGxE is not as restrictive as one might naively assume. In an MR context there are limited cases in which a confounder will be a determinant of a genetic instrument, though this could occur through mechanisms such as assortative mating. This is only problematic, however, if the confounder simultaneously associated with the interaction-covariate. It seems most likely that MRGxE estimates will exhibit bias where the instrument is a determinant of one or more confounders, which in turn are determinants of the interaction covariate. We therefore recommend care be taken in examining such pathways and suggest the use of MRGxE as a sensitivity analysis for detecting pleiotropic bias.

### MRGxE as a sensitivity analysis

In cases where the constant pleiotropy assumption is assumed to be violated, MRGxE can still be used in sensitivity analyses as a means to select a subset of valid instruments. To show how this is the case, we begin by clarifying that an invalid instrument can be detected in principle whenever 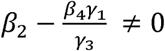, due to either *β*_2_ ≠ 0, *β*_4_ ≠ 0, or both. As a consequence, MRGxE can be used to assess the validity of individual instruments, informing instrument selection and components of allelic scores. There are, however, two important considerations when applying this approach. First, it is not possible to distinguish the average pleiotropic effect across interaction-covariate subgroups from the change in pleiotropic effect between instrument-covariate subgroups. It is therefore a test of invalidity occurring either due to an average non-zero pleiotropic effect across interaction-covariate subgroups, or due to changing pleiotropic effects between interaction-covariate subgroups, and cannot be used to correct MRGxE estimates directly.

Second, MRGxE will incorrectly fail to detect invalid instruments in the special case where:

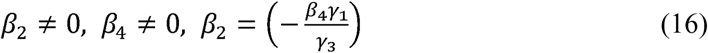

Given that this second scenario is very unlikely, it is possible to still utilise MRGxE when assumptions are violated as a sensitivity analysis for assessing whether a directional pleiotropic effect is present, however, it would not be possible to produce an unbiased association estimate.

### Causal effect of BMI upon SBP

There exists an extensive literature on the relationship between adiposity and SBP, with both observational(34) and MR(35–37) studies finding evidence of positive association. However, the magnitude of this association has been found to differ markedly between such studies, with observational studies often recording greater effect sizes than those using MR.

As an applied example, we perform two sample summary MR and MRGxE analyses examining the effect of adiposity (measured using BMI) upon SBP using data from the GIANT consortium(29) and UK Biobank. The motivation in performing both forms of analysis is to highlight the extent to which pleiotropic effect estimates obtained using MRGxE with single instrument agree with conventional MR approaches. In the first instance analyses are performed using the full set of participants for which data is available, after which analyses were repeated restricting the analyses to 334976 non-related participants of European ancestry.

In conducting a two-sample summary analysis, effect estimates and standard errors for 95 genetic variants associated with BMI (*p* = 5 × 10^−8^) were obtained from Locke et al(29) using the GIANT consortium sample (with a full list of variants presented in the web appendix). Corresponding estimates for each genetic variant with respect to SBP were obtained using UK Biobank. In contrast, MRGxE was implemented by constructing a weighted allelic score using estimates from the GIANT consortium. The MRGxE analysis can be viewed as analogous to two-sample summary data MR, using instrument-exposure estimates for BMI as external weights, and individual data from a separate sample to inform instrument-outcome association estimates. In each analysis BMI, SBP, and the weighted allelic score were standardised.

### Analysis I: Two-Sample Summary Analysis

We implement several two-sample summary MR methods utilising the mrrobust software package(38) in Stata SE 14.0(39). Performing IVW provides an estimate comparable to TSLS, and produces estimates with greater precision than alternative summary approaches. However, as IVW estimates can exhibit bias in the presence of horizontal pleiotropy, MR-Egger regression, weighted median, and weighted modal approaches are also considered as sensitivity analyses.

A range of methods are adopted in sensitivity analyses for two key reasons. First, each method relies upon differing assumptions with respect to the underlying distribution of pleiotropic effects. MR-Egger regression requires the effect of genetic variants on the exposure to be independent of their pleiotropic effects on the outcome (InSIDE)(5). The weighted median requires more than 50% of variants (with respect to their weighting) to be valid instruments(10), whilst the modal estimator assumes that the most frequent value of the pleiotropic bias across the set of genetic variants is zero (ZEMPA) (40).

Estimates for each method using the unrestricted UK Biobank sample are presented in Table 1, with an accompanying plot showing the IVW and MR-Egger estimates in Figure 4.

**Table 1:** A table showing two sample summary MR estimates for the effect of BMI upon SBP using the unrestricted UK Biobank sample. A smoothing parameter (*ϕ* = 1) was selected in implementing the modal estimator, and a value 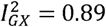 using MR Egger is indicative of regression dilution of approximately 11% towards the null. MR Egger estimates using SIMEX to correct for regression dilution are also presented.

| Method | Estimate | SE | 95% CI | p.value |
| --- | --- | --- | --- | --- |
| IVW | 0.103 | 0.030 | (0.04, 0.16) | <0.001 |
| MR-Egger (intercept) | 0.003 | 0.002 | (-0.00, 0.01) | 0.212 |
| MR-Egger (effect)<sup>1</sup> | 0.014 | 0.077 | (-0.14, 0.16) | 0.851 |
| SIMEX corrected MR-Egger (intercept) | 0.003 | 0.003 | (-0.00, 0.01) | 0.380 |
| SIMEX corrected MR-Egger (effect) | 0.018 | 0.121 | (-0.22, 0.25) | 0.880 |
| Weighted Median | 0.130 | 0.020 | (0.09, 0.17) | <0.001 |
| Modal Estimator<sup>2</sup> | 0.133 | 0.027 | (0.08, 0.19) | <0.001 |

<sup>1</sup> $I_{GX}^2 = 0.88$ <sup>2</sup> Smoothing parameter $\phi = 1$

**Figure 4.**
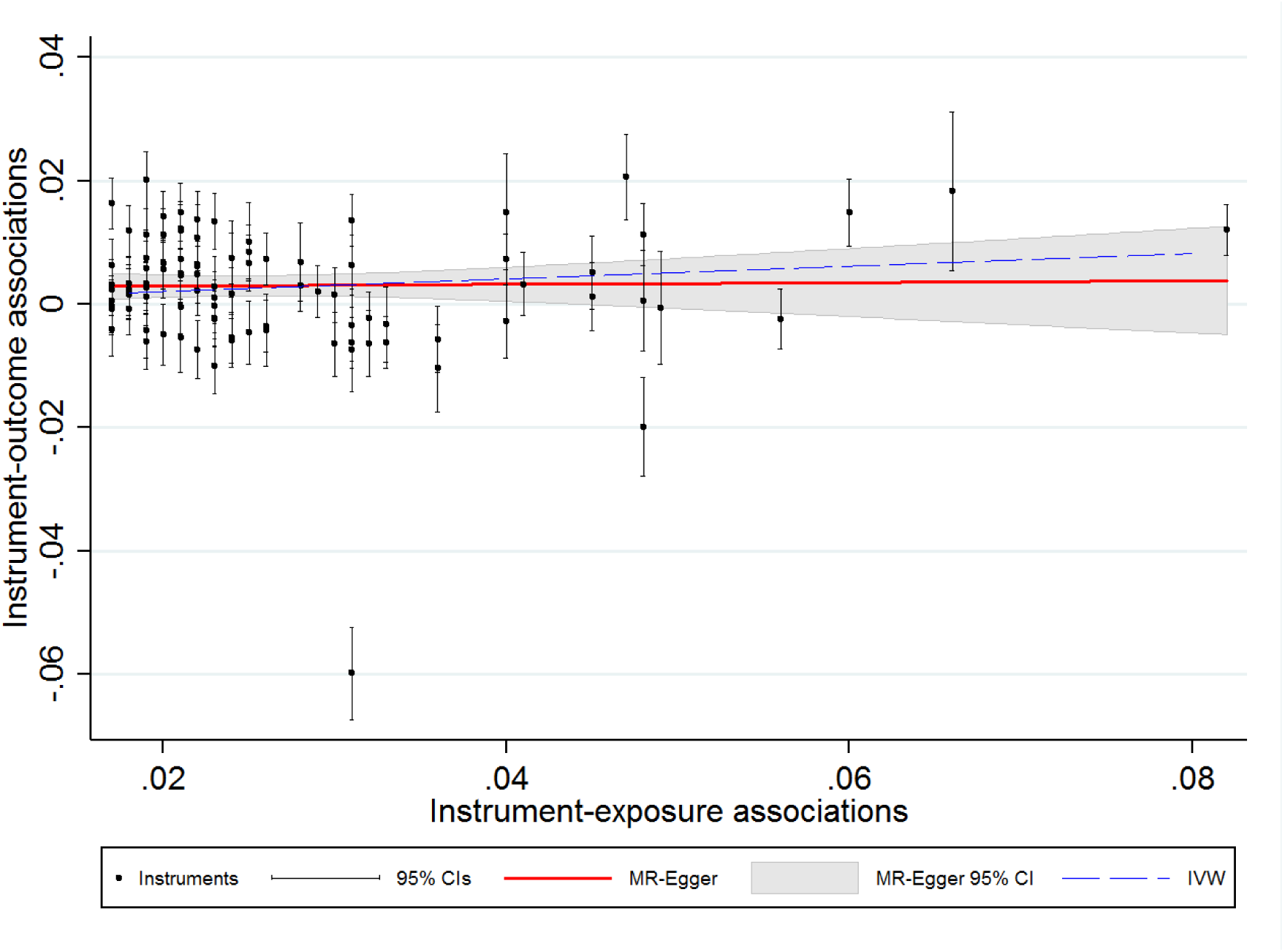
Scatter plot showing IVW and MR-Egger estimates for the effect of BMI upon SBP using unrestricted UK Biobank sample

With the exception of MR-Egger regression, each of the methods performed above show evidence of a positive association between BMI and SBP. There does not appear to be evidence of a substantial unbalanced horizontal pleiotropic effect using either MR-Egger, weighted median, or weighted modal approaches, with the IVW estimate lying within the confidence intervals of both the weighted median and weighted modal estimates.

Disagreement in effect estimation between MR-Egger regression and the other methods can in part be explained by regression dilution bias, as evidenced by an 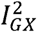 value of 89.5 (indicative of a relative bias of 10.5% towards the null), as well as by identifying influential outliers. We apply a SIMEX correction for regression dilution bias following Bowden et al(41), though the inference remains consistent with the unadjusted MR-Egger estimates. Outliers can be identified by calculating the degree of heterogeneity utilising Rucker’s Q statistic, and estimating the relative contribution of each genetic variant to overall heterogeneity, as outlined in Bowden et al(41). In doing so, the overall Rucker’s *Q* statistic is 1069.42 (*p* =< 0.001), with rs11191560 contributing almost a quarter of the overall heterogeneity (*Q* = 274.4, *p* = < 0.001).

Estimates for the restricted UK Biobank sample are presented in Table 2, with an accompanying plot showing the IVW and MR-Egger estimates in Figure 5.

**Table 2:**
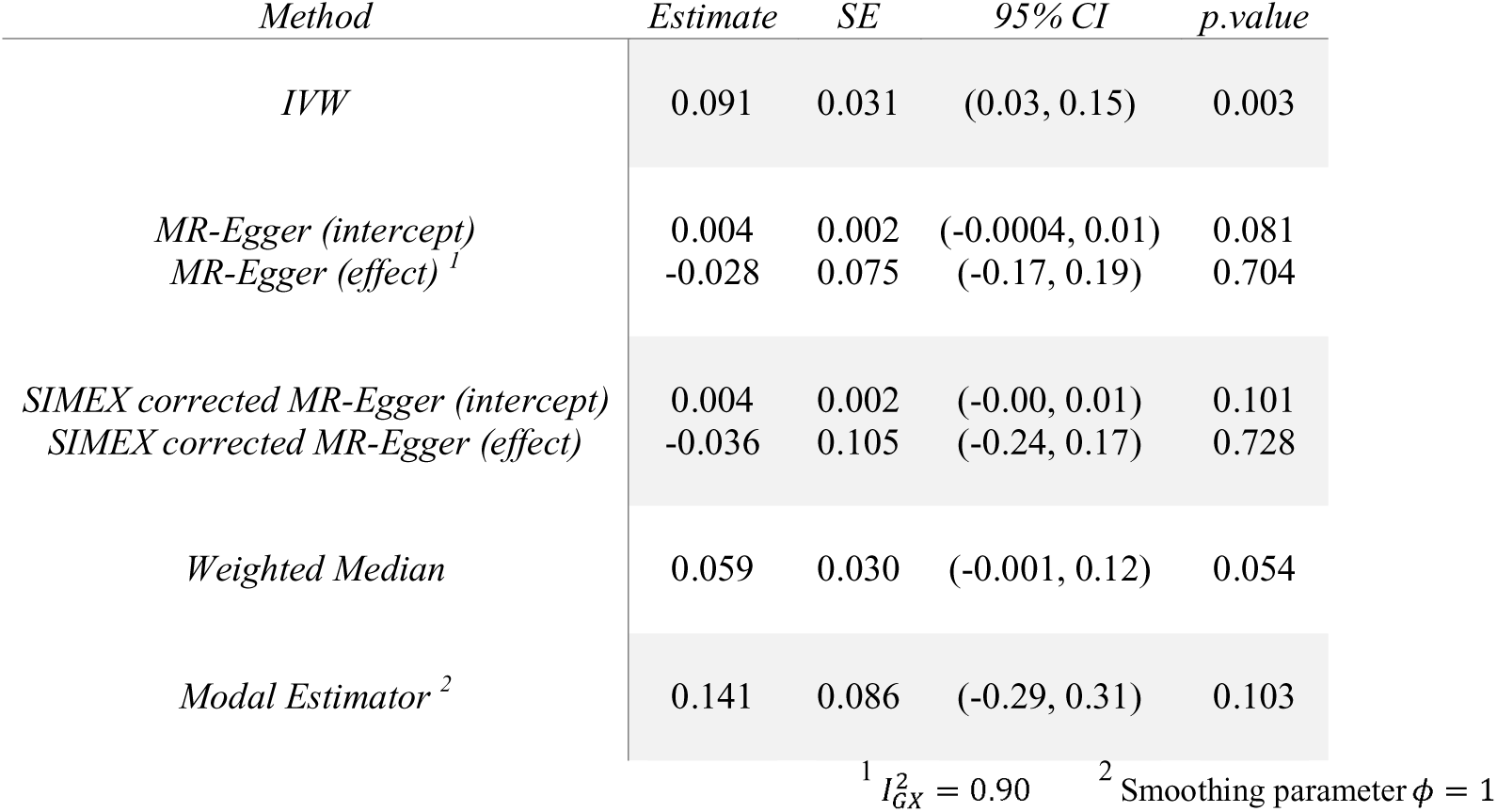
A table showing two sample summary MR estimates for the effect of BMI upon SBP using the restricted UK Biobank sample. A smoothing parameter (*ϕ* = 1) was selected in implementing the modal estimator, and a value 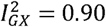 using MR Egger is indicative of regression dilution of approximately 10% towards the null.

**Figure 5.**
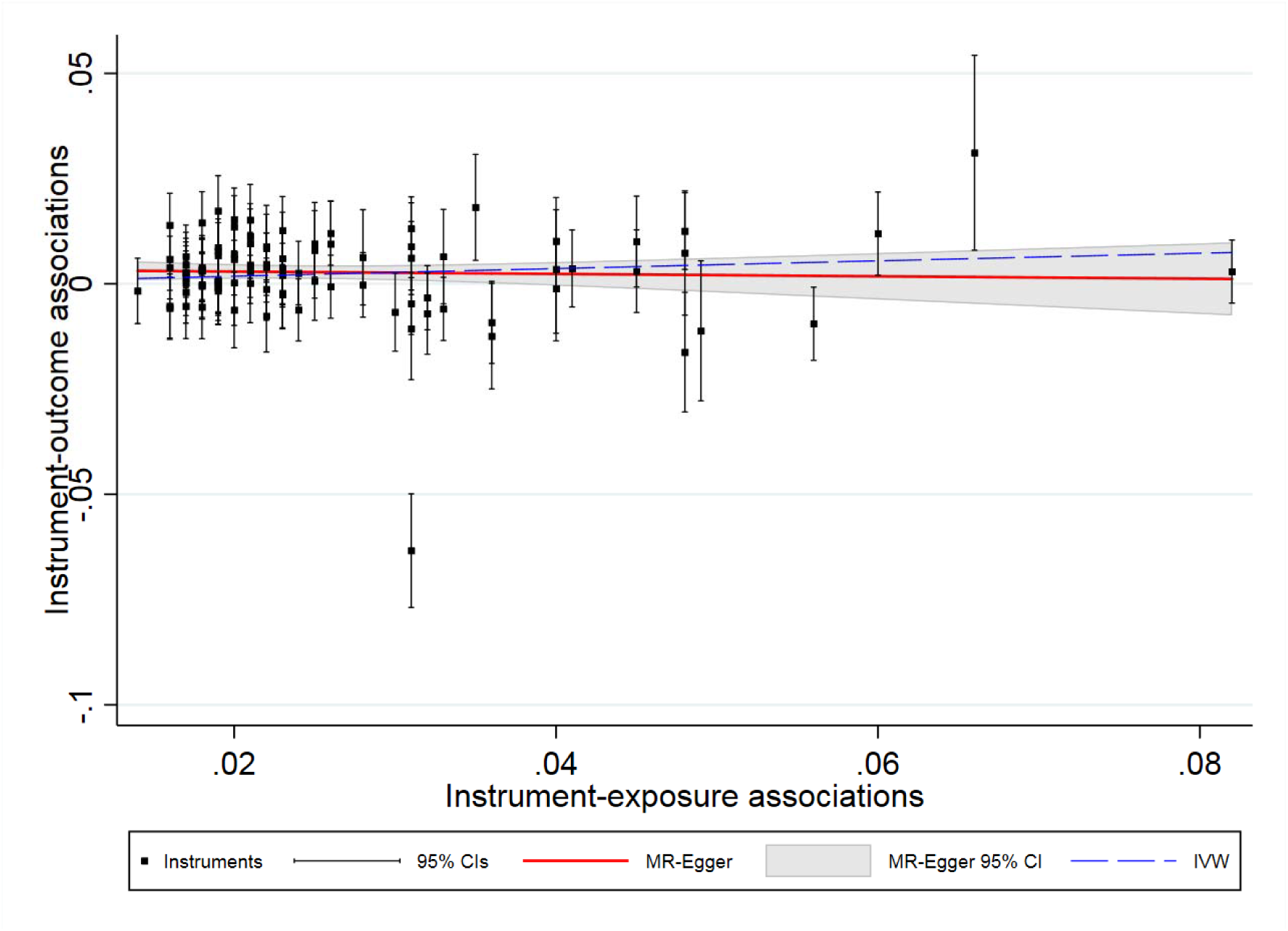
Scatter plot showing IVW and MR-Egger estimates for the effect of BMI upon SBP using restricted UK Biobank sample.

When excluding related participants and individuals with non-European ancestry sensitivity analyses indicate the presence of pleiotropic bias. There are at least two possible explanations for the disagreement in estimates between the unrestricted and restricted analyses. Initially, including participants of differing ancestries can introduce bias into MR estimates, as the characteristics of the genetic instruments (for example in terms of frequency) can be markedly different between population groups. However, by excluding individuals on the basis of ancestry we may inadvertently also be conditioning on further variables outside of ethnicity and relatedness, consequently inducing selection bias. For the purpose of this analysis, the primary goal is to assess the extent to which MR Egger and MRGxE produce estimates which are in agreement, serving as motivation for presenting MRGxE analyses using unrestricted and restricted samples from UK Biobank for comparison with the MR Egger estimates.

### Analysis II: MRGxE using Townsend Deprivation Index

In implementing MRGxE, Townsend Deprivation Index (TDI) was selected as a continuous covariate for which instrument strength was expected to vary, based on findings from previous studies(23, 42). TDI is a common derived measure of socio-economic deprivation, using many variables such as car ownership, occupation type and educational attainment(43).

In the UK Biobank, TDI scores were obtained from preceding national census data and calculated for electoral districts *(“wards”* comprised of approximately 5,500 individuals). Participants were assigned a score based upon the area in which they lived, determined using the postcode of their home dwelling. The selection of TDI was based upon previous evidence suggesting genetically determined BMI to be a weaker predictor of BMI for individuals experiencing lower levels of social deprivation(23). Missing values were considered to be missing completely at random (MCAR), and were removed prior to performing the analysis. This resulted in a total of 193106 participants with complete data.

We present observational and TSLS estimates using the weighted allelic score as an instrument and controlling for TDI in tables 3 and 4 for the unrestricted and restricted samples respectively. In both cases, we find evidence of a positive association between BMI and SBP, with a greater magnitude of effect for the observational estimate.

**Table 3:** OLS and TSLS effect estimates for the unrestricted UK Biobank sample. The OLS estimates are obtained by regressing SBP upon BMI and TDI, whilst the TSLS estimates utilise the weighted allelic score as an instrument for BMI.

|  | *Estimate* | *SE* | *95% CI* | *p.value* |
| --- | --- | --- | --- | --- |
| **OLS** |  |  |  |  |
| *Intercept* | <0.0001 | 0.002 | (-0.004, 0.004) | >0.999 |
| *BMI* | 0.192 | 0.002 | (0.19, 0.20) | <0.001 |
| *TDI* | -0.056 | 0.002 | (-0.06, -0.05) | <0.001 |
| **TSLS** |  |  |  |  |
| *Intercept* | <0.0001 | 0.002 | (-0.004, 0.004) | >0.999 |
| *BMI* | 0.129 | 0.014 | (0.10, 0.16) | <0.001 |
| *TDI* | -0.050 | 0.002 | (-0.054, -0.047) | <0.001 |

**Table 4:** OLS and TSLS effect estimates for the restricted UK Biobank sample. The OLS estimates are obtained by regressing SBP upon BMI and TDI, whilst the TSLS estimates utilise the weighted allelic score as an instrument for BMI.

|  | *Estimate* | *SE* | *95% CI* | *p.value* |
| --- | --- | --- | --- | --- |
| **OLS** |  |  |  |  |
| *Intercept* | <0.001 | 0.002 | (-0.004, 0.004) | >0.999 |
| *BMI* | 0.190 | 0.002 | (0.185, 0.194) | <0.001 |
| *TDI* | -0.048 | 0.002 | (-0.05, -0.04) | <0.001 |
| **TSLS** |  |  |  |  |
| *Intercept* | <0.001 | 0.002 | (-0.004, 0.004) | >0.999 |
| *BMI* | 0.105 | 0.017 | (0.07, 0.14) | <0.001 |
| *TDI* | -0.040 | 0.003 | (-0.05, -0.04) | <0.001 |

For both the unrestricted and restricted samples the estimates agree with findings of the previous studies discussed above. The instrument is also considered to be sufficiently strong to overcome weak instrument bias, with an F statistic of 3623 in the restricted sample. To perform MRGxE, we divided the sample on the basis of TDI score into 5, 10, 20, and 50 population subgroups. In each case, a ratio estimate was calculated for each group, after which IVW and MRGxE estimates were produced. The results of each analysis are presented in tables 5 and 6, with IVW referring to an inverse-variance weighted estimate using interaction covariate subgroups.

**Table 5:** IVW and MRGxE Estimates for different numbers of TDI quantile groupings using the unrestricted UK Biobank sample For each group, the IVW estimate represents an inverse weighted estimate using each of the TDI subgroups, providing an estimate equivalent to the TSLS estimates using the weighted allelic score.

| *Number of groups* | *Method* | *Estimate* | *SE* | *95% CI* | *p.value* |
| --- | --- | --- | --- | --- | --- |
| 5 | MRGxE (intercept) | 0.002 | 0.005 | (-0.02, 0.02) | 0.728 |
|  | MRGxE (effect) | 0.114 | 0.040 | (-0.02, 0.24) | 0.065 |
|  | IVW | 0.129 | 0.004 | (0.12, 0.14) | <0.001 |
| 10 | MRGxE (intercept) | 0.006 | 0.013 | (-0.02, 0.04) | 0.670 |
|  | MRGxE (effect) | 0.087 | 0.096 | (-0.01, 0.31) | 0.391 |
|  | IVW | 0.129 | 0.012 | (0.10, 0.16) | <0.001 |
| 20 | MRGxE (intercept) | 0.002 | 0.015 | (-0.03, 0.03) | 0.915 |
|  | MRGxE (effect) | 0.117 | 0.110 | (-0.11, 0.35) | 0.300 |
|  | IVW | 0.129 | 0.015 | (0.10, 0.16) | <0.001 |
| 50 | MRGxE (intercept) | -0.008 | 0.013 | (-0.03, 0.02) | 0.514 |
|  | MRGxE (effect) | 0.190 | 0.094 | (0.00, 0.38) | 0.050 |
|  | IVW | 0.129 | 0.012 | (0.10, 0.16) | <0.001 |

**Table 6:** IVW and MRGxE Estimates for different numbers of TDI quantile groupings using the restricted UK Biobank sample For each group, the IVW estimate represents an inverse weighted estimate using each of the TDI subgroups, providing an estimate equivalent to the TSLS estimates using the weighted allelic score.

| *Number of groups* | *Method* | *Estimate* | *SE* | *95% CI* | *p.value* |
| --- | --- | --- | --- | --- | --- |
| 5 | MRGxE (intercept) | 0.011 | 0.009 | (-0.02, 0.04) | 0.298 |
|  | MRGxE (effect) | 0.020 | 0.066 | (-0.19, 0.23) | 0.779 |
|  | IVW | 0.102 | 0.010 | (0.07, 0.13) | <0.001 |
| 10 | MRGxE (intercept) | 0.013 | 0.009 | (-0.01, 0.03) | 0.191 |
|  | MRGxE (effect) | 0.004 | 0.068 | (-0.15, 0.16) | 0.952 |
|  | IVW | 0.101 | 0.012 | (0.08, 0.12) | <0.001 |
| 20 | MRGxE (intercept) | 0.017 | 0.011 | (-0.01, 0.04) | 0.144 |
|  | MRGxE (effect) | -0.025 | 0.083 | (-0.20, 0.15) | 0.765 |
|  | IVW | 0.100 | 0.013 | (0.07, 0.13) | <0.001 |
| 50 | MRGxE (intercept) | 0.001 | 0.013 | (-0.03, 0.03) | 0.924 |
|  | MRGxE (effect) | 0.094 | 0.097 | (-0.10, 0.29) | 0.340 |
|  | IVW | 0.103 | 0.017 | (0.07, 0.14) | <0.001 |

From Table 5 we see that the IVW estimates are directionally consistent and of a similar magnitude to the TSLS estimates as expected. In each case, there again appears to be limited evidence of pleiotropy, whilst there appears to be some indication of a positive effect of BMI upon SBP, particularly in the 5-group case. Figure 6 displays both the IVW and MRGxE estimates for the 5-group case, whilst corresponding plots for other groups are presented in the web appendix.

**Figure 6:**
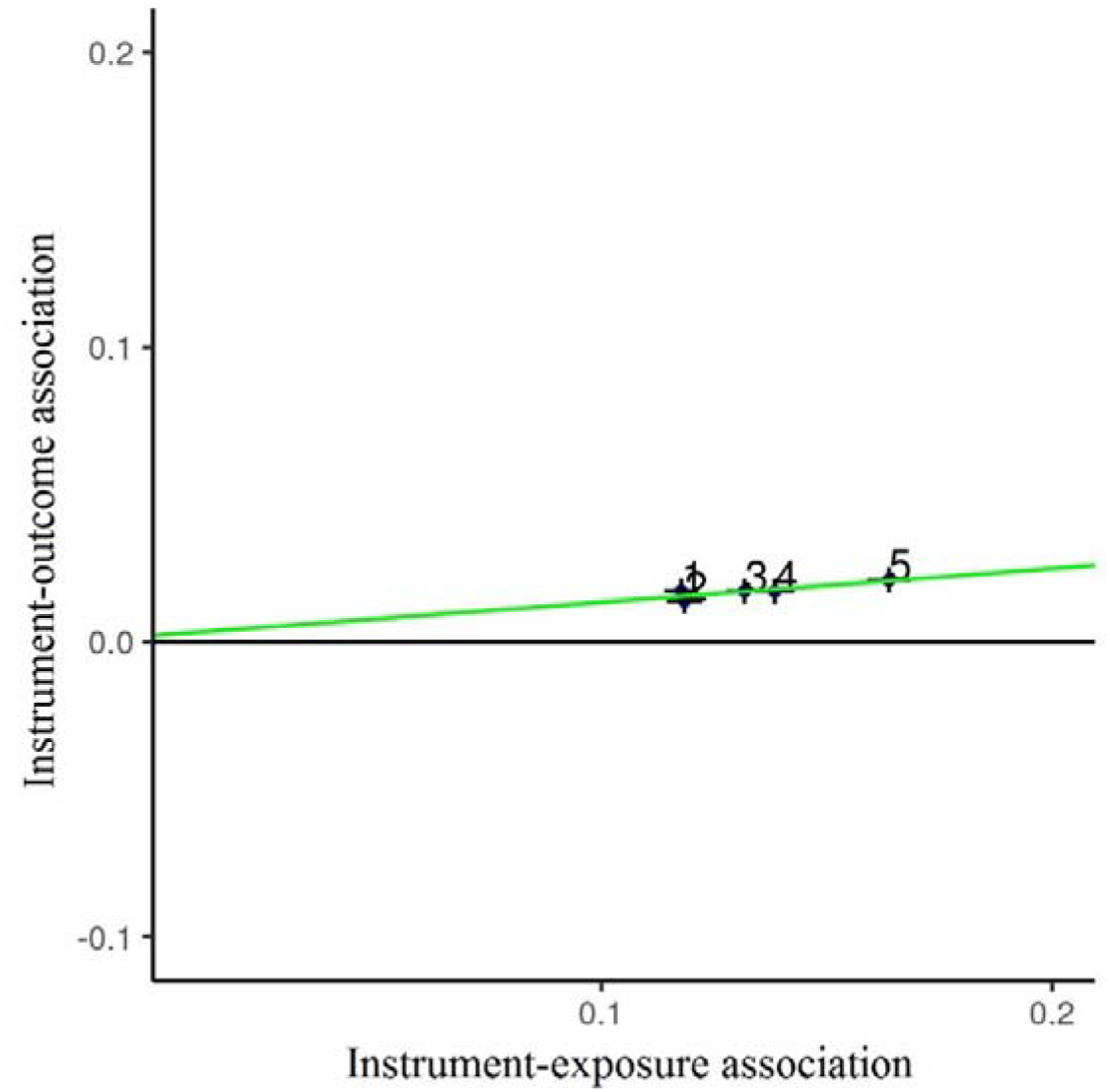
A scatterplot showing the MRGxE estimate (indicated in green) for the unrestricted UK Biobank sample. Each point represents ascending quintiles of Townsend Deprivation Index, in this case showing the strength of the instrument-exposure association to increase with increasing socio-economic deprivation. The close proximity of the intercept to the origin is indicative of instrument validity.

Considering Figure 6, a number of key features of the analysis can be identified. Initially, the ordering of the TDI groups supports the assumption that the instrument-exposure association varies across levels of TDI. In particular, the least deprived groups (group 1 and group 2) have the weakest association, suggesting that genetically predicted BMI is a weaker predictor of BMI for participants experiencing lower levels of deprivation. A further observation is that the positioning of each estimate provides some evidence of a linear interaction, with instrument strength increasing monotonically as subgroup TDI increases.

Table 6 shows the MRGxE estimates corresponding to the restricted UK Biobank sample across the same range of interaction-covariate groupings. As with the unrestricted sample, the estimates are in agreement with two sample summary estimates. Figure 7 shows a plot corresponding to 5 group MRGxE analysis using the restricted sample. Unlike the unrestricted sample analysis, there appears to be evidence of bunching in the lower TDI groups, suggesting non-linearity in the interaction term.

**Figure 7:**
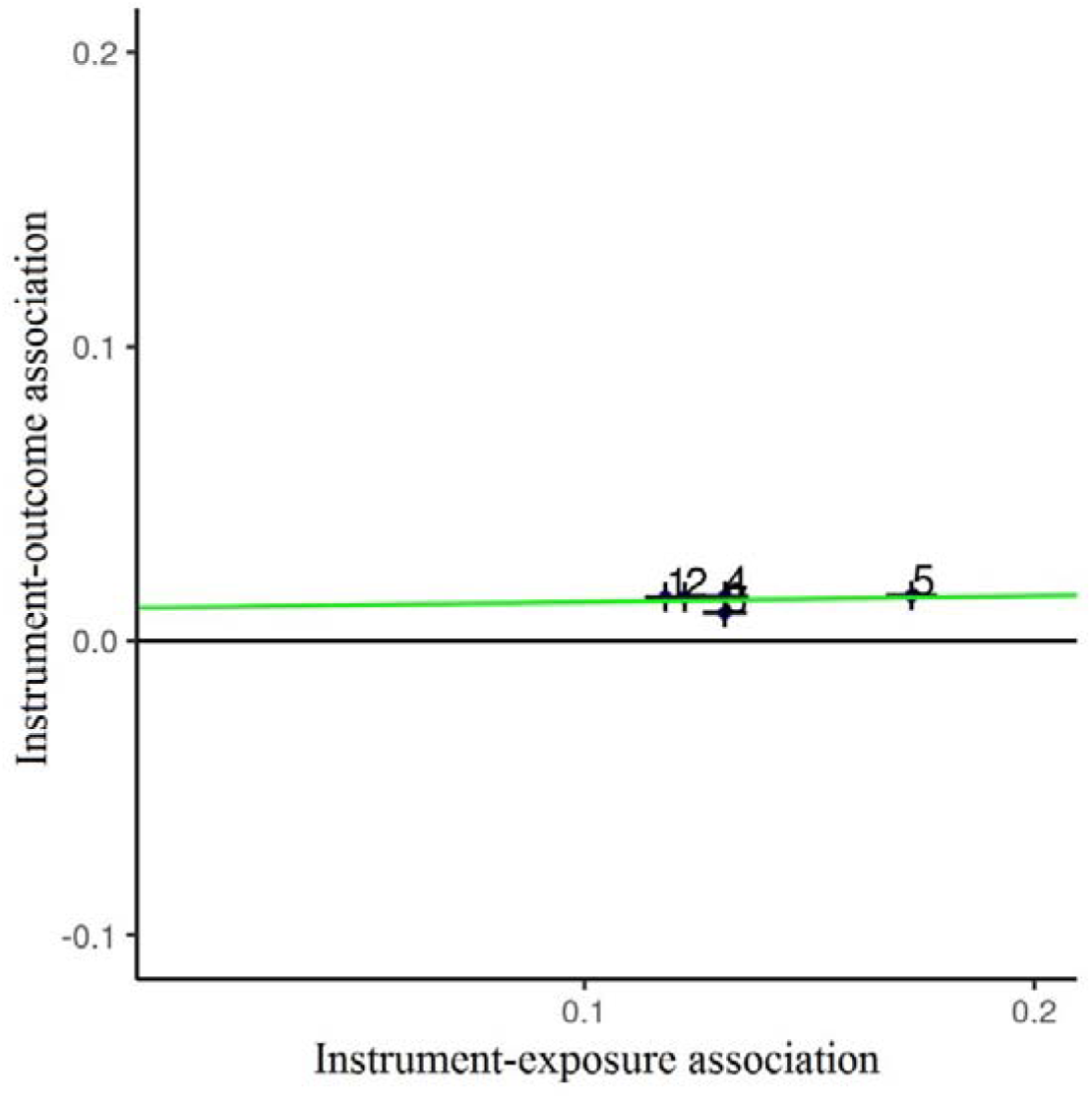
A scatterplot showing the MRGxE estimate (indicated in green) for the restricted UK Biobank sample. Each point represents ascending quintiles of Townsend Deprivation Index, in this case showing the strength of the instrument-exposure association to increase with increasing socio-economic deprivation. Contrary to Figure 6, the deviation of the intercept from the origin is indicative of instrument invalid instrument use.

Comparing these estimates to those obtained using two-sample summary MR, there appears to be substantial agreement in the findings of the two approaches. MRGxE provides estimates of horizontal pleiotropy and causal effect equivalent to MR Egger in the summary MR context, and constraining the MRGxE model to the intercept yields an effect estimate similar to IVW. The lack of agreement in detecting a positive effect of BMI upon SBP using conventional thresholds can be attributed in part to insufficient strength of TDI as in interacting covariate. The observed agreement in the findings between the single instrument and multiple instrument approaches is encouraging, and similar results are produced using alternate interaction-covariates such as frequency of alcohol consumption (shown in the web appendix).

One important consideration in performing MR analyses is that causal effect estimates are often uncertain, due to either a lack of precision or doubts regarding the assumptions of the implemented approach. A response to this issue put forward by VanderWeele et al(44), has been to shift the emphasis from identifying the magnitude of causal effects to identifying the presence of causal effects. Under such a paradigm, estimation using instrument-covariate interactions, such as through MRGxE, can be particularly insightful in identifying broad effects or associations in epidemiological studies. Adopting this rationale, MRGxE can be used as a broad test of instrument validity in cases where the underlying assumptions of the approach are likely violated, focusing on effect direction as opposed to the magnitude of the estimated effect.

### Simulations

To illustrate the effectiveness of MRGxE, and further consider the importance of the constant pleiotropy assumption with respect to causal effect estimation, we perform a simulation study within a two-sample MR framework. Considering a realistic case, two sets of simulations are performed, the first using a null causal effect (*β*_1_ = 0), and the second a positive causal effect (*β*_1_ = 0.05). Individual level data is generated, from which the necessary summary data estimates are extracted. In each case, a total of 5 population subgroups are considered, with further details provided in the web appendix.

Four distinct cases are considered:

- No pleiotropy and the constant pleiotropy assumption satisfied
- Directional pleiotropy and the constant pleiotropy assumption satisfied
- No pleiotropy and the constant pleiotropy assumption violated
- Directional pleiotropy and the constant pleiotropy assumption violated

The results for each case represent the mean values for 10 000 simulated datasets.

## Results

Results of the simulation analysis are presented in Table 7 and Table 8 representing the null effect and 0.05 causal effect scenarios respectively. The mean F statistic remains the same for each case, with substantial variation in F statistic between interaction covariate groups. This is essential, as the variation in instrument strength can be viewed as variation in instrument relevance for particular population subgroups. In this case, estimates using IVW and MRGxE, as well as significance values were taken directly from each regression output without using regression weights, as the variant-outcome associations were found to have the same standard errors.

**Table 7:** Performance of IVW and MRGxE methods in simulation setting with null causal effect *β*_1_ = 0. In case 4, *β*_4_ = *β*_2_ and consequentially the intercept of the MRGxE model is approximately 0. This explains the similarity in the power of the MRGxE pleiotropy test between Case 1 and Case 4.

| *Case* | *N* | *N Per Decile* | *Mean F Statistic* | *IVW*<br>*Mean Estimate*<br>*(mean SE)* | *IVW*<br>*Type I Error*<br>*Rate* | *MRGxE*<br>*Mean Estimate*<br>*(mean SE)* | *MRGxE*<br>*Power of*<br>*Pleiotropy Test* | *MRGxE*<br>*Effect*<br>*Type I Error*<br>*Rate* |
| --- | --- | --- | --- | --- | --- | --- | --- | --- |
| *Case 1:*<br>$\beta_1 = 0$<br>$\beta_2 = 0$<br>$\beta_4 = 0$ | 10000 | 2000 | 80.1 | 0.003 (0.046) | 0.049 | 0.004 (0.066) | 0.047 | 0.050 |
|  | 20000 | 4000 | 159.3 | 0.002 (0.033) | 0.050 | 0.003 (0.047) | 0.048 | 0.050 |
|  | 30000 | 6000 | 238.3 | 0.001 (0.027) | 0.052 | 0.001 (0.038) | 0.050 | 0.050 |
|  | 40000 | 8000 | 317.3 | 0.001 (0.023) | 0.052 | 0.001 (0.033) | 0.049 | 0.049 |
|  | 50000 | 10000 | 396.2 | 0.000 (0.021) | 0.050 | 0.001 (0.030) | 0.049 | 0.052 |
| *Case 2:*<br>$\beta_1 = 0$<br>$\beta_2 = 0.05$<br>$\beta_4 = 0$ | 10000 | 2000 | 80.1 | 0.091 (0.060) | 0.153 | 0.004 (0.066) | 0.209 | 0.050 |
|  | 20000 | 4000 | 159.3 | 0.089 (0.051) | 0.168 | 0.003 (0.047) | 0.365 | 0.050 |
|  | 30000 | 6000 | 238.3 | 0.089 (0.048) | 0.157 | 0.001 (0.038) | 0.504 | 0.050 |
|  | 40000 | 8000 | 317.3 | 0.088 (0.047) | 0.150 | 0.001 (0.033) | 0.612 | 0.049 |
|  | 50000 | 10000 | 396.2 | 0.088 (0.046) | 0.135 | 0.001 (0.030) | 0.708 | 0.052 |
| *Case 3:*<br>$\beta_1 = 0$<br>$\beta_2 = 0$<br>$\beta_4 = 0.05$ | 10000 | 2000 | 80.1 | 0.082 (0.059) | 0.130 | 0.169 (0.061) | 0.262 | 0.429 |
|  | 20000 | 4000 | 159.3 | 0.081 (0.051) | 0.128 | 0.169 (0.043) | 0.442 | 0.673 |
|  | 30000 | 6000 | 238.3 | 0.080 (0.048) | 0.121 | 0.167 (0.035) | 0.579 | 0.816 |
|  | 40000 | 8000 | 317.3 | 0.080 (0.047) | 0.100 | 0.167 (0.031) | 0.676 | 0.895 |
|  | 50000 | 10000 | 396.2 | 0.079 (0.046) | 0.091 | 0.167 (0.027) | 0.770 | 0.945 |
| *Case 4:*<br>$\beta_1 = 0$<br>$\beta_2 = 0.05$<br>$\beta_4 = 0.05$ | 10000 | 2000 | 80.1 | 0.169 (0.043) | 0.783 | 0.169 (0.061) | 0.048 | 0.429 |
|  | 20000 | 4000 | 159.3 | 0.168 (0.030) | 0.962 | 0.169 (0.043) | 0.051 | 0.673 |
|  | 30000 | 6000 | 238.3 | 0.167 (0.025) | 0.994 | 0.167 (0.035) | 0.051 | 0.816 |
|  | 40000 | 8000 | 317.3 | 0.167 (0.022) | 0.999 | 0.167 (0.031) | 0.048 | 0.895 |
|  | 50000 | 10000 | 396.2 | 0.167 (0.019) | 1.000 | 0.167 (0.027) | 0.047 | 0.945 |

**Table 8:** Performance of IVW and MRGxE methods in simulation setting with positive causal effect *β*_1_ = 0.05. In case 8, *β*_4_ = *β*_2_ and consequentially the intercept of the MRGxE model is approximately 0. This explains the similarity in the power of the MRGxE pleiotropy test between Case 5 and Case 8.

| *Case* | *N* | *N Per Decile* | *Mean F Statistic* | *IVW*<br>*Mean Estimate*<br>*(mean SE)* | *IVW*<br>*Power to detect*<br>*causal effect* | *MRGxE*<br>*Mean Estimate*<br>*(mean SE)* | *MRGxE*<br>*Power of*<br>*pleiotropy Test* | *MRGxE*<br>*Power to detect*<br>*causal effect* |
| --- | --- | --- | --- | --- | --- | --- | --- | --- |
| *Case 5:*<br>$\beta_1 = 0.05$<br>$\beta_2 = 0$<br>$\beta_4 = 0$ | 10000 | 2000 | 80.1 | 0.053 (0.046) | 0.145 | 0.054 (0.066) | 0.047 | 0.093 |
|  | 20000 | 4000 | 159.3 | 0.052 (0.033) | 0.219 | 0.053 (0.047) | 0.048 | 0.123 |
|  | 30000 | 6000 | 238.3 | 0.051 (0.027) | 0.288 | 0.051 (0.038) | 0.050 | 0.148 |
|  | 40000 | 8000 | 317.3 | 0.051 (0.023) | 0.355 | 0.049 (0.033) | 0.049 | 0.181 |
|  | 50000 | 10000 | 396.2 | 0.050 (0.021) | 0.423 | 0.051 (0.030) | 0.049 | 0.207 |
| *Case 6:*<br>$\beta_1 = 0.05$<br>$\beta_2 = 0.05$<br>$\beta_4 = 0$ | 10000 | 2000 | 80.1 | 0.141 (0.060) | 0.381 | 0.054 (0.066) | 0.209 | 0.093 |
|  | 20000 | 4000 | 159.3 | 0.139 (0.051) | 0.495 | 0.053 (0.047) | 0.365 | 0.123 |
|  | 30000 | 6000 | 238.3 | 0.139 (0.048) | 0.553 | 0.051 (0.038) | 0.504 | 0.146 |
|  | 40000 | 8000 | 317.3 | 0.138 (0.047) | 0.597 | 0.051 (0.033) | 0.612 | 0.181 |
|  | 50000 | 10000 | 396.2 | 0.138 (0.046) | 0.630 | 0.051 (0.030) | 0.708 | 0.207 |
| *Case 7:*<br>$\beta_1 = 0.05$<br>$\beta_2 = 0$<br>$\beta_4 = 0.05$ | 10000 | 2000 | 80.1 | 0.132 (0.059) | 0.341 | 0.219 (0.061) | 0.262 | 0.615 |
|  | 20000 | 4000 | 159.3 | 0.131 (0.051) | 0.417 | 0.219 (0.043) | 0.442 | 0.854 |
|  | 30000 | 6000 | 238.3 | 0.130 (0.048) | 0.474 | 0.217 (0.035) | 0.579 | 0.945 |
|  | 40000 | 8000 | 317.3 | 0.129 (0.047) | 0.510 | 0.217 (0.031) | 0.676 | 0.981 |
|  | 50000 | 10000 | 396.2 | 0.129 (0.046) | 0.534 | 0.217 (0.027) | 0.770 | 0.993 |
| *Case 8:*<br>$\beta_1 = 0.05$<br>$\beta_2 = 0.05$<br>$\beta_4 = 0.05$ | 10000 | 2000 | 80.1 | 0.219 (0.043) | 0.933 | 0.219 (0.061) | 0.048 | 0.615 |
|  | 20000 | 4000 | 159.3 | 0.218 (0.030) | 0.997 | 0.219 (0.043) | 0.051 | 0.854 |
|  | 30000 | 6000 | 238.3 | 0.217 (0.025) | 1.000 | 0.217 (0.035) | 0.051 | 0.945 |
|  | 40000 | 8000 | 317.3 | 0.217 (0.022) | 1.000 | 0.217 (0.031) | 0.048 | 0.981 |
|  | 50000 | 10000 | 396.2 | 0.217 (0.019) | 1.000 | 0.217 (0.027) | 0.047 | 0.993 |

Initially, cases within the null causal effect scenario are considered. In the valid instrument case, both IVW and MRGxE provide unbiased causal effect estimates, though the IVW estimate is more accurate. This is similar to comparisons between IVW and MR-Egger regression, supporting use of IVW in cases where directional pleiotropy is absent. Type I error rates remained at approximately 5% for both IVW and MRGxE, and in testing for directional pleiotropy. In the second case, estimates using IVW are biased in the direction of pleiotropic association, whilst MRGxE continues to produce unbiased estimates. This bias appears to decrease marginally with sample size increases. As the sample size increases, power to detect directional pleiotropy using MRGxE increases from 21% to 70%, whilst Type I error rates remain at a nominal 5% level.

The third case represents a situation in which the instrument is not valid, but the degree of pleiotropy changes between population subgroups. In this case, both IVW and MRGxE produce biased causal effect estimates, though the MRGxE causal effect estimates exhibit a greater degree of bias than the IVW estimates. This contributes to an increase in Type I error rate relative to IVW, which estimates the causal effect to be smaller in magnitude. In this situation, the MRGxE test for directional pleiotropy is particularly powerful, rising from 43% to 95% as the sample size increases. This seeming increase in power can be attributed violation of the constant pleiotropy assumption (*β*_4_ ≠ 0) leading to biased pleiotropy estimates (*β*_2_), in this case overestimating the magnitude of pleiotropic effects. In the final case, both IVW and MRGxE produce estimates with similar sizes of bias and precision. A particularly interesting feature of this case is that the MRGxE test for directional pleiotropy is suggestive of a null pleiotropic effect, remaining at 5%. This represents a situation in which *β*_2_ = *β*_4_, invalidating the use of MRGxE as a sensitivity analysis. In the positive causal effect scenario, both the MRGxE and IVW approaches produce estimates exhibiting similar patterns to those in the null causal effect case.

## Discussion

In this paper, we have presented a method to identify and correct for pleiotropic bias in MR studies using instrument-covariate interactions. In cases where the constant pleiotropy assumption is satisfied, individual instruments can be assessed, providing less biased causal effect estimates compared to conventional estimates such as IVW in the presence of directional pleiotropy. Where individual level data are available, and where it is sensible to assume an underlying linear model, the Cho et al(21) approach is appropriate and provides estimates in agreement with MRGxE.

However, in cases where directional pleiotropy is not present, IVW is a more accurate method and should be preferred. In cases where the constant pleiotropy assumption is violated, a sensible approach would be to prune invalid variants using the pleiotropy estimates from MRGxE, and then implement IVW using the set of valid variants. In this sense, MRGxE can be viewed as a sensitivity analysis in a similar fashion to MR-Egger, which can be applied to a single genetic instrument within the individual level data setting(5, 45).

### Comparison with existing methods

MRGxE represents a synthesis of both Slichter regression(14) and MR-Egger regression. As with Slichter regression, the method can be applied outside of an MR context, provided that an interaction can be identified which induces variation in instrument strength, and conforms with the interaction restrictions previously discussed. In Slichter 2014, an example of assessing returns to schooling conditioning on IQ levels represents such an application(14, 16). The similarities with MR-Egger regression are such that, rather than a competing methodology, it is more correctly viewed as a counterpart to the method.

It is likely that median based approaches can be applied within the MRGxE framework, and this forms an promising avenue for future study. In the summary data setting, such methods rely upon at least 50% of the instruments being valid, or 50% of the instruments being valid with respect to their weighting when implementing weighted median regression. An analogous approach within the MRGxE framework would require at least 50% of the population subgroups to satisfy the constant pleiotropy assumption, as these have a similar role to individual instruments in the summary setting.

Finally PRMR, a closely related approach to detecting and correcting for pleiotropy has recently been proposed by van Kippersluis and Rietveld(46). Under this framework, in cases where a no relevance group is observed, the degree of association between the instrument and the outcome is equated with the exact pleiotropic effect across the whole population. In this respect the approach is similar to that of Chen et al(26). This term is then incorporated as an offset within a standard analysis. Whilst their approach is useful in highlighting the potential of no relevance groups in assessing pleiotropy, it can be criticised for ignoring true uncertainty in the pleiotropic effect estimate. Its practical application is also limited by the fact that strict instrument-exposure independence is rare. For example, the authors cite Cho et al’s(21) MR analysis using gender specific alcohol consumption as a canonical example(46), but in fact 25% of female participants in this study did consume alcohol(21). Such a violation would obviously undermine an approach that assumed a strict no relevance group. This serves as motivation for the development of a formal statistical model (MRGxE) to use variation in gene-exposure associations across a covariate to infer the likely location of a no relevance group whilst properly accounting for its uncertainty, and use this as a basis for detecting and adjusting for pleiotropy.

### Two-Sample Summary MRGxE

Whilst this paper has focused primarily on the application of MRGxE to individual level data (albeit by extracting and then meta-analysing summary statistics obtained from it), it clearly applies to cases where subgroup specific summary data on instrument-exposure and instrument-outcome associations are available. An alternative approach would be to meta-analyse summary statistics obtained from many separate studies under the assumption that study-specific estimates relate to a study-specific characteristic. For example, the work of Robinson et al(32) highlights the interaction between age and adult BMI heritability as one potential candidate, given that age is likely to vary naturally across contributing studies.

### Limitations of MRGxE

There are a number of factors which must be considered before implementing MRGxE. Firstly, the constant pleiotropy assumption is essential for causal estimate correction. If there is reason to believe that pleiotropic effects differ between population subgroups, then using the approach will result in misleading causal effect estimates. One useful aspect to this problem, however, is that provided the first stage interaction is sufficiently strong, bias from changes in pleiotropic effect may be sufficiently small as to be negligible in analyses. This may well be the case in situations such as the Cho et al(21) study, where the difference in instrument effect between gender groups is very strong in comparison to potential variation in pleiotropic effect. As it is not possible to directly measure the change in pleiotropic effect across groups, decisions regarding appropriate instrument-covariate interaction selection require justification.

A second limitation of the approach is that, owing to the limited availability of summary data estimates for particular covariate groups, it may be difficult to implement in a summary data setting. At present researchers may be limited to common groupings such as gender, unless further information is made available upon request. A second complication using the summary MRGxE approach focuses upon the use of study heterogeneity. In many cases, the degree to which such heterogeneity is present with respect to the instrument-covariate interaction may be insufficient to perform a meaningful analysis. A related concern is that studies exhibiting such heterogeneity may undermine the extent to which homogeneity in remaining effects can be assumed. This can introduce confounding and undermine subsequent inference.

A further consideration pertaining to the majority of methods, including MRGxE, is the extent to which the study sample is representative of the sample of interest. In cases where the sample is not representative, selection bias can have a substantial impact on resulting estimates. This is illustrated in the difference between the estimated effect of BMI upon SBP using the interim UK Biobank release, in which the inclusion of a disproportionate number of heavy smokers may have resulted in a 3-fold increase in the magnitude of the estimated effect (as given in the web appendix). As a number of previous studies have found evidence suggesting an interaction between genetically predicted BMI and smoking(42, 47), this could highlight smoking as a potential confounder violating the interaction-covariate restrictions outlined in this paper. It may also be possible that composite phenotypes (such as scores derived from several measures) may have differing contributions to the outcome at differing interaction-covariate levels.

This limitation can somewhat be mitigated by performing multiple iterations of MRGxE using a set of interaction covariates. Provided that the instrument-covariate interaction of sufficient strength, it would be expected that resulting estimates would be in agreement. In cases where substantial disagreement is observed, such disagreement could be indicative of violation of the constant pleiotropy assumption, or characteristics of the underlying confounding structure. The works of Emdin et al(22) and Krishna et al(48) follow this reasoning. Further work will consider the implications of interaction-covariate selection, and role of confounding within the context of MRGxE.

Finally, it is important to consider results from MR gene-environment interaction approaches within the context of existing evidence using alternate approaches. This underpins the concept of triangulation put forward by Lawlor, Tilling, and Davey Smith(49), in which differences in estimates across a range of approaches can be indicative of sources of bias potentially unique to each research design. Identifying disagreement in estimated effects across studies of differing design can therefore prove valuable in identifying avenues for further research, whilst substantial agreement strengthens confidence in the resulting findings and subsequent inference.

## Conclusion

This paper formalises an intuitive method for assessing pleiotropic bias, which has gained increasing traction in recent years. At present, MRGxE serves as a valuable test for directional pleiotropy, and can provide causal effect estimates robust to directional pleiotropy in cases where the constant pleiotropy assumption is satisfied. It is therefore most appropriate for use as a sensitivity analysis in studies using individual level data, and in informing instrument selection and allelic score construction.

## Supplementary Material

A web appendix containing supplementary materials can be found at:

## Acknowledgements

The authors would like to thank Gibran Hemani and James Staley for assistance in data formatting, as well as Yoonsu Cho, Neil Davies, Fernando Hartwig, and Tom Palmer for useful comments and suggestions on this work.

This research has been conducted using the UK Biobank Resource under Application Number 8786.

## References

1. Davey Smith G, Ebrahim S. ‘Mendelian randomization’: can genetic epidemiology contribute to understanding environmental determinants of disease? Int J Epidemiol. 2003;32(1):1–22.

2. Davey Smith G, Hemani G. Mendelian randomization: genetic anchors for causal inference in epidemiological studies. Hum Mol Genet. 2014;23(R1):R89–98.

3. Burgess S, Small DS, Thompson SG. A review of instrumental variable estimators for Mendelian randomization. Statistical Methods in Medical Research. 2015:0962280215597579.

4. Glymour MM, Tchetgen Tchetgen EJ, Robins JM. Credible Mendelian randomization studies: approaches for evaluating the instrumental variable assumptions. Am J Epidemiol. 2012;175(4):332–9.

5. Bowden J, Davey Smith G, Burgess S. Mendelian randomization with invalid instruments: effect estimation and bias detection through Egger regression. Int J Epidemiol. 2015;44(2):512–25.

6. Burgess S, Thompson SG. Multivariable Mendelian randomization: the use of pleiotropic genetic variants to estimate causal effects. Am J Epidemiol. 2015;181(4):251–60.

7. Burgess S, Thompson SG. Use of allele scores as instrumental variables for Mendelian randomization. Int J Epidemiol. 2013;42(4):1134–44.

8. Davies NM, von Hinke Kessler Scholder S, Farbmacher H, Burgess S, Windmeijer F, Davey Smith G. The many weak instruments problem and Mendelian randomization. Stat Med. 2015;34(3):454–68.

9. Burgess S, Butterworth A, Thompson SG. Mendelian randomization analysis with multiple genetic variants using summarized data. Genet Epidemiol. 2013;37(7):658–65.

10. Bowden J, Davey Smith G, Haycock PC, Burgess S. Consistent Estimation in Mendelian Randomization with Some Invalid Instruments Using a Weighted Median Estimator. Genet Epidemiol. 2016;40(4):304–14.

11. Hartwig FP, Davey Smith G, Bowden J. Robust inference in summary data Mendelian randomization via the zero modal pleiotropy assumption. Int J Epidemiol. 2017;46(6):1985–98.

12. Hartwig FP, Davies NM. Why internal weights should be avoided (not only) in MR-Egger regression. Int J Epidemiol. 2016;45(5):1676–8.

13. Bowden J, Burgess S, Davey Smith G. Response to Hartwig and Davies. Int J Epidemiol. 2016;45(5):1679–80.

14. Slichter D. Testing Instrument Validity and Identification with Invalid Instruments. University of Rochester: Rochester: USA; 2014.

15. Imbens GW, Angrist JD. Identification and Estimation of Local Average Treatment Effects. Econometrica. 1994;62(2):467–75.

16. Card D. Using Geographic Variation in College Proximity to Estimate the Return to Schooling. National Bureau of Economic Research, Inc; 1993.

17. Conley TG, Hansen CB, Rossi PE. Plausibly Exogenous. Rev Econ Stat. 2012;94(1):260–72.

18. Gennetian L, Bos, J.M, Morris, P.A. Using Instrumental Variables Analysis to Learn More from Social Policy Experiments. MDRC Working Papers on Research Methodology. 2002.

19. Small DS. Mediation analysis without sequential ignorability: Using baseline covariates interacted with random assignment as instrumental variables. Journal of Statistical Research. 2012.

20. van Kippersluis H, Rietveld CA. Pleiotropy-robust Mendelian randomization. Int J Epidemiol. 2017.

21. Cho Y, Shin SY, Won S, Relton CL, Davey Smith G, Shin MJ. Alcohol intake and cardiovascular risk factors: A Mendelian randomisation study. Scientific Reports. 2015;5.

22. Emdin CA, Khera AV, Natarajan P, Klarin D, Zekavat SM, Hsiao AJ, et al. Genetic Association of Waist-to-Hip Ratio With Cardiometabolic Traits, Type 2 Diabetes, and Coronary Heart Disease. Jama-Journal of the American Medical Association. 2017;317(6):626–34.

23. Tyrrell J, Yaghootkar H, Beaumont R, Jones SE, Ames RM, Tuke MA, et al. Gene-obesogenic environment interactions in the UK Biobank study. Diabetologia. 2016;59:S51–S.

24. Davey Smith G. Use of genetic markers and gene-diet interactions for interrogating population-level causal influences of diet on health. Genes and Nutrition. 2011;6(1):27–43.

25. Lewis SJ, Davey Smith G. Alcohol, ALDH2, and esophageal cancer: A meta-analysis which illustrates the potentials and limitations of a Mendelian randomization approach. Cancer Epidemiology Biomarkers & Prevention. 2005;14(8):1967–71.

26. Chen L, Davey Smith G, Harbord RM, Lewis SJ. Alcohol intake and blood pressure: A systematic review implementing a Mendelian Randomization approach. Plos Medicine. 2008;5(3):461–71.

27. Davey Smith G. Mendelian Randomization for Strengthening Causal Inference in Observational Studies: Application to Gene x Environment Interactions. Perspectives on Psychological Science. 2010;5(5):527–45.

28. Bulik-Sullivan BK, Loh PR, Finucane HK, Ripke S, Yang J, Patterson N, et al. LD Score regression distinguishes confounding from polygenicity in genome-wide association studies. Nature Genetics. 2015;47(3):291-+.

29. Locke AE, Kahali B, Berndt SI, Justice AE, Pers TH, Day FR, et al. Genetic studies of body mass index yield new insights for obesity biology. Nature. 2015;518(7538):197–206.

30. Taylor AE, Lu F, Carslake D, Hu Z, Qian Y, Liu S, et al. Exploring causal associations of alcohol with cardiovascular and metabolic risk factors in a Chinese population using Mendelian randomization analysis. Sci Rep. 2015;5:14005.

31. Freathy RM, Kazeem GR, Morris RW, Johnson PC, Paternoster L, Ebrahim S, et al. Genetic variation at CHRNA5-CHRNA3-CHRNB4 interacts with smoking status to influence body mass index. Int J Epidemiol. 2011;40(6):1617–28.

32. Robinson MR, English G, Moser G, Lloyd-Jones LR, Triplett MA, Zhu Z, et al. Genotype-covariate interaction effects and the heritability of adult body mass index. Nat Genet. 2017;49(8):1174–81.

33. Wald A. The Fitting of Straight Lines if Both Variables are Subject to Error. 1940:284–300.

34. Droyvold WB, Midthjell K, Nilsen TI, Holmen J. Change in body mass index and its impact on blood pressure: a prospective population study. Int J Obes (Lond). 2005;29(6):650–5.

35. Li LJ, Liao J, Cheung CY, Ikram MK, Shyong TE, Wong TY, et al. Assessing the Causality between Blood Pressure and Retinal Vascular Caliber through Mendelian Randomisation. Sci Rep. 2016;6:22031.

36. Timpson NJ, Harbord R, Davey Smith G, Zacho J, Tybjaerg-Hansen A, Nordestgaard BG. Does greater adiposity increase blood pressure and hypertension risk?: Mendelian randomization using the FTO/MC4R genotype. Hypertension. 2009;54(1):84–90.

37. Holmes MV, Lange LA, Palmer T, Lanktree MB, North KE, Almoguera B, et al. Causal effects of body mass index on cardiometabolic traits and events: a Mendelian randomization analysis. Am J Hum Genet. 2014;94(2):198–208.

38. Spiller W, Davies NM, Palmer TM. Software Application Profile: mrrobust - A Tool For Performing Two-Sample Summary Mendelian Randomization Analyses. bioRxiv. 2017.

39. StataCorp. Stata Statistical Software: Release 14. College Station, TX: StataCorp LP. 2015.

40. Hartwig FP, Davey Smith G, Bowden J. Robust inference in summary data Mendelian randomization via the zero modal pleiotropy assumption. International Journal of Epidemiology. 2017.

41. Bowden J, Del Greco M F, Minelli C, Lawlor D, Sheehan N, Thompson J, et al. Improving the accuracy of two-sample summary data Mendelian randomization: moving beyond the NOME assumption. bioRxiv. 2017.

42. Rask-Andersen M, Karlsson T, Ek WE, Johansson Å. Gene-environment interaction study for BMI reveals interactions between genetic factors and physical activity, alcohol consumption and socioeconomic status. PLOS Genetics. 2017;13(9):e1006977.

43. Mackenbach JP. Health and Deprivation - Inequality and the North - Townsend,P, Phillimore,P, Beattie,A. Health Policy. 1988;10(2):207-.

44. VanderWeele TJ, Tchetgen Tchetgen EJ, Cornelis M, Kraft P. Methodological challenges in mendelian randomization. Epidemiology. 2014;25(3):427–35.

45. Lewis BC, Nair PC, Heran SS, Somogyi AA, Bowden JJ, Doogue MP, et al. Warfarin resistance associated with genetic polymorphism of VKORC1: linking clinical response to molecular mechanism using computational modeling. Pharmacogenet Genomics. 2016;26(1):44–50.

46. van Kippersluis H, Rietveld CA. Pleiotropy-robust Mendelian Randomization. bioRxiv. 2016.

47. Justice AE, Winkler TW, Feitosa MF, Graff M, Fisher VA, Young K, et al. Genome-wide meta-analysis of 241,258 adults accounting for smoking behaviour identifies novel loci for obesity traits. Nat Commun. 2017;8:14977.

48. Krishna A, Razak F, Lebel A, Davey Smith G, Subramanian SV. Trends in group inequalities and interindividual inequalities in BMI in the United States, 1993-2012. Am J Clin Nutr. 2015;101(3):598–605.

49. Lawlor DA, Tilling K, Davey Smith G. Triangulation in aetiological epidemiology. International Journal of Epidemiology. 2016;45(6):1866–86.

